# Shearing in flow environment promotes evolution of social behavior in microbial populations

**DOI:** 10.1101/198507

**Authors:** Gurdip Uppal, Dervis Can Vural

## Abstract

It is advantageous for microbes to form social aggregates when they commonly benefit from secreting a public good. However, cooperating microbial groups can be evolutionarily unstable, since a cheating strain that does not secrete the public good can reproduce quicker and take over. Here we study the effects of fluid advection patterns on group reproduction as a mechanism to enable or enhance social behavior in microbial populations. We use a realistic advection-diffusion-reaction model to describe microbial growth and mutation in a flow environment. Social groups arise naturally from our model as self-reproducing Turing patterns that can avoid mutant takeovers at steady state. Our central finding is that flow shear enables and promotes social behavior in microbes by limiting the spread of cheating strains. Regions of the flow domain with higher shear admits high cooperativity and large population density, whereas low shear regions are devoid of life due to opportunistic mutations.

## Introduction

Cooperation is the cement of biological complexity. A combined investment brings larger returns. However, while cooperating groups are fitter, individuals in these groups have evolutionary incentive to cheat by taking advantage of available public goods without contributing their own. Avoiding the cost of these goods allow larger reproduction rates, causing cheaters to proliferate until the lack of public goods compromise the fitness of the entire group. In other words, while cooperating groups are fitter than non–cooperating ones, cooperation is not evolutionarily stable. How then, can social behavior emerge and persist in microbial colonies?

The evolution of cooperation is an active field of research, with multiple theories resolving this dilemma ***Axelrod and Hamilton*** (***1981***); ***Sachs et al.*** (***2004***); ***Sachs and Simms*** (***2006***); ***Nowak*** (***2006***). These can be grouped into two broad categories. Theories in the first category argue that persistent cooperation is a direct or indirect byproduct of a self-serving trait, so that “altruism is not really altruistic” (or conversely, “selfishness is not really selfish”). For example, the metabolic byproducts of one organism may be a resource for another, and vice versa. Another example is positive ***Trivers*** (***1971***) and negative ***Clutton-Brock and Parker*** (***1995***); ***El Mouden et al.*** (***2010***) reciprocity, where cooperators are rewarded later by others, or where cheaters are inflicted a cost, via policing or reputation. For example, quorum signals reveal whether available public goods add up to the population density. In this case, altruists cut back public good production to eliminate cheaters (albeit with collateral damage) ***Allen et al.*** (***2016***); ***Sandoz et al.*** (***2007***); ***Diggle et al.*** (***2007***).

The second class of theories employ classical selection arguments at larger scales or sub-populations ***Grafen*** (***1984***). Group selection ***Wynne-Edwards*** (***1962***); ***Haldane*** (***1932***); ***Traulsen and Nowak*** (***2006***); ***Wilson*** (***1975***) and its modern incarnation, multi-level selection, ***Wilson and Sober*** (***1994***) propose that cooperating groups (or groups of groups) will reproduce faster than non-cooperating ones and prevail. A closely related idea, kin-selection ***Hamilton*** (***1964a***,b); ***Williams*** (***1966***); ***Smith*** (***1964***); ***Hamilton*** (***1975***); ***Lion et al.*** (***2011***), proposes that individuals cooperate with those to which they are genetically related, and thus, a cooperative genotype is really cooperating with itself.

Hamilton conjectured that kin selection should promote cooperation if the population is viscous, i.e. when the mobility of the population is limited ***Hamilton*** (***1964a***,b). This helps ensure that genetically related individuals cooperate with each other. However, competition within kin can inhibit altruism ***Taylor*** (***1992***); ***Wilson et al.*** (***1992***). One solution to this is if individuals disperse as groups, also known as budding dispersal. This was shown to promote cooperation theoretically by ***Gardner and West*** (***2006***) and demonstrated experimentally by ***Kümmerli et al.*** (***2009a***). The physical mechanism offered in the present theoretical study may be seen as a way of inducing or enhancing budding dispersal in planktonic microbial systems.

Typically, evolution of cooperation is quantitatively analyzed with the aid of game theoretic models applied to well-mixed populations, networks and other phenomenological spatial structures ***Szabó and Fath*** (***2007***); ***Allen et al.*** (***2013***); ***Nowak and Sigmund*** (***2004***); ***Vural et al.*** (***2015***). While there are few models that take into account spatial proximity effects, ***Medvinsky et al.*** (***2002***); ***Nadell et al.*** (***2010***, 2013); ***Dobay et al.*** (***2014***); ***Driscoll and Pepper*** (***2010***) and the influence of decay and diffusion of public goods ***Dobay et al.*** (***2014***); ***Wakano et al.*** (***2009***); ***Hauert et al.*** (***2008***), how advective fluid flow influences social evolution remains unexplored.

A flowing habitat can have a drastic effect on population dynamics ***Tel et al.*** (***2005***); ***Nickerson et al.*** (***2004***); ***Koshel’ and Prants*** (***2006***); ***Sandulescu et al.*** (***2008***). For example, a flowing open system can allow the coexistence of species despite their differential fitness ***Karolyi et al.*** (***2000***). Interactions between fluid shear and bacterial motility has been shown to lead to shear trapping ***Rusconi et al.*** (***2014***); ***Berke et al.*** (***2008***) which causes preferential attachment to surfaces ***Berke et al.*** (***2008***); ***Li et al.*** (***2011***). Turbulent flows can also lead to a trade-off in nutrient uptake and the cost of locomotion due to chemotaxis ***Taylor and Stocker*** (***2012***), and can drastically effect the population density ***Pigolotti et al.*** (***2012***); ***Perlekar et al.*** (***2010***). Most importantly, the reproductive successes of species (and individuals within a single species) are coupled over distance, through the secretion of toxins, goods, and signals ***Mimura et al.*** (***2000***); ***Allison*** (***2005***); ***Hibbing et al.*** (***2010***). The spatial distributions of all such fittness altering intermediaries depend on flow, and thus, we are motivated to find out how flow plays a role in the evolution of cooperation.

Here we theoretically study how fluid dynamics molds the sociality of a population of planktonic species. Qualitatively stated, our evolutionary model has three assumptions: (1) Individuals secrete one waste compound and one public good. The former has no cost, whereas the latter does. (2) Mutations can vary the public good secretion rates of microbes, thereby producing a continuum of social behavior. (3) Microbes and their secretions diffuse and flow according to the laws of fluid dynamics.

Thus, our model is applicable to a wide variety of social ecosystems ranging from phytoplankton flowing in oceanic currents to opportunistic bacteria colonizing blood or industrial pipelines. Our findings imply that that greater social complexity amongst planktonic species can be observed in regions of large shear, such as by rocks and river banks. We might even speculate that multicellularity may have originated near fluid domains with large shear flow, rather than the bulk of oceans or lakes.

This paper is organized as follows: We first establish that under certain conditions our physical model gives rise to spatially organized cooperative groups. The groups are a natural byproduct of the convection-diffusion dynamics of the system. Furthermore, under certain physical conditions, we show that these social structures reproduce in whole to form new identical structures, essentially giving rise to a “group selection” setting. We then study the effects of mutation, and observe that above a certain mutation rate cheating strains take over groups which leads to extinction. The latter finding is consistent with other empiric ***Rainey and Rainey*** (***2003***); ***Diggle et al.*** (***2007***) and theoretical studies ***Nowak and May*** (***1992***).

After setting up the stage for naturally forming social groups, we present our central finding, that flow shear promotes and enhances cooperativity within these groups. Specifically, we demonstrate and study the evolution of sociality of a microbial population (1) subjected to constant shear, (2) embedded in a cylindrical laminar flow and (3) in a Rankine vortex. We find that in all three cases population density and cooperative behavior scales with flow shear. The mechanism of action is that shear distortion fragments cooperative clusters, thereby limiting the spread of cheaters. If the shear is large enough that groups are torn apart at a larger rate than the mutation rate, then cooperation will prevail.

## Methods

We study a realistic spatial model of microorganisms, where fluid dynamical forces contribute significantly to the evolution of their social behavior. Our analysis consists of simulations and analytical formulas. It is well known that evolution of altruism in species strongly depends on the individuals being discrete ***Durrett and Levin*** (***1994***). We therefore simulate microbes as discrete particles subject to stochastic physical and evolutionary forces. We describe compounds secreted by microbes as continuous fields. In contrast, our analytical expressions are derived from analogous equations that are entirely continuous and deterministic. We compare the outcomes of our agent-based simulation with analytical formulas based on continuous equations.

In our model, the microorganisms secrete two types of diffusive molecules that influence each others fitness (Fig.1). The first molecule, the concentration of which denoted by *c*_1_(*x, t*), is a beneficial public good that increases the fitness of those exposed, whereas the second, *c*_2_(*x, t*), is a waste compound or toxin that has the opposite effect. The continuous equations that represent our system are

**Figure 1.**
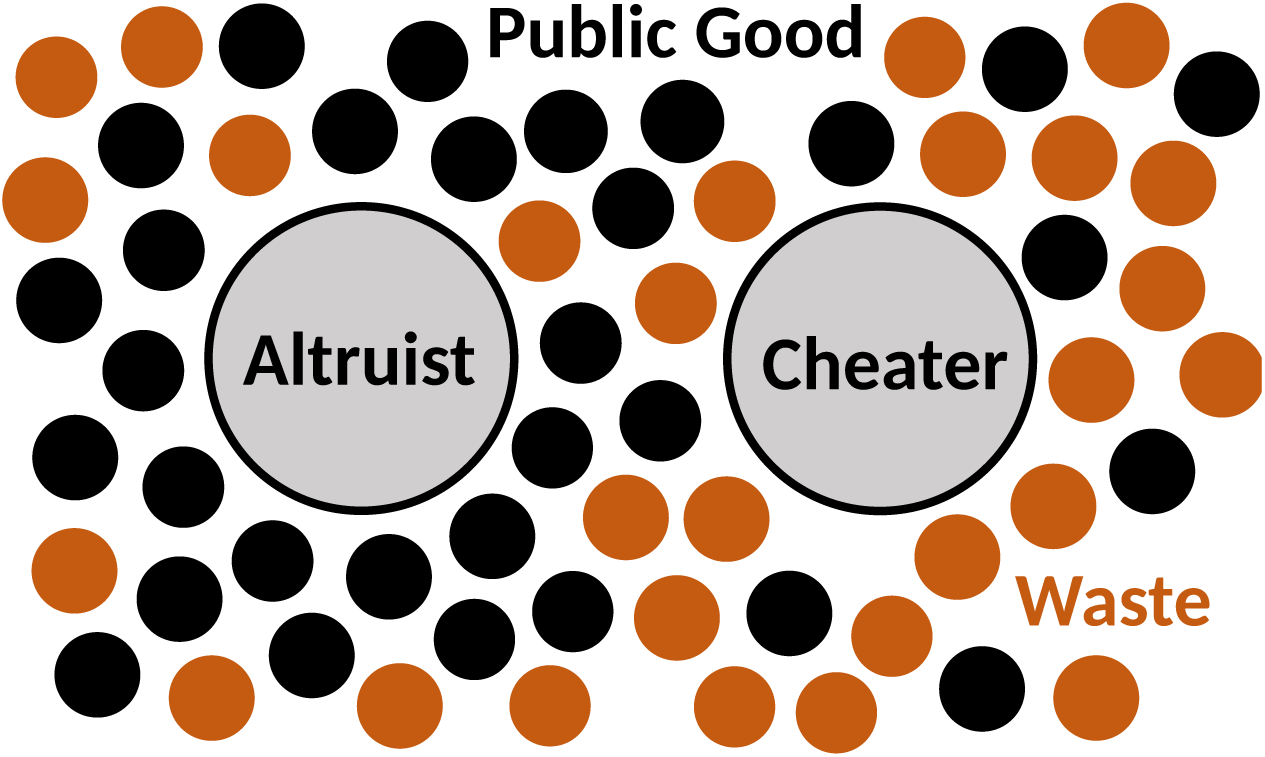
Schematics of our Model. The microbes secrete two types of molecules into the environment. The first, a beneficial public good that promotes growth, and the second, a waste or harmful substance hinders growth. Cheating microbes produce less or none of the former, while benefiting from public goods secreted by the cooperating population.

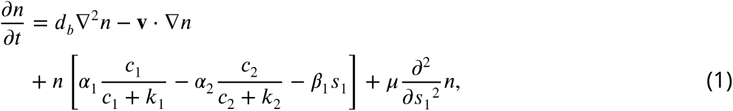

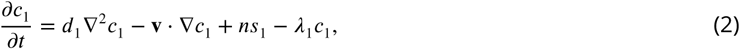

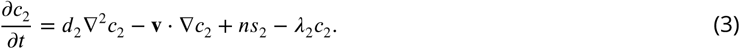

Here *n* is a shorthand for *n*(*x, t, s*_1_), the number density of microbes at time *t* and position *x* that produce the public good at a rate of *s*_1_. These microbes pay a fitness cost of *β*_1_*s*_1_ per unit time. The production rate of waste *s*_2_ on the other hand, is assumed constant for all, and has no fitness cost. Waste limits the number of individuals that a unit volume can carry. Microbes secreting public goods at a rate *s*_1_ replicate to produce others with the same secretion rate. This reproduction rate is given by the square bracket. However, the production rate *s*_1_ can change due to mutations. This is described by the last term of (1). Mutations can be thought as diffusion in *s*_1_ space.

In all three equations the first two terms describe diffusion and advection, while the last two terms of (2) and (3) describe the production and decay of chemicals. The first two terms in the square bracket describe the effect of the secreted compounds on fitness. This saturating form is experimentally established and well understood ***Monod*** (***1949***). The crucial third term in the square bracket describes the cost of producing the public good.

Through some rescaling, we can non-dimensionalize the system. If we define the rescaled variables as

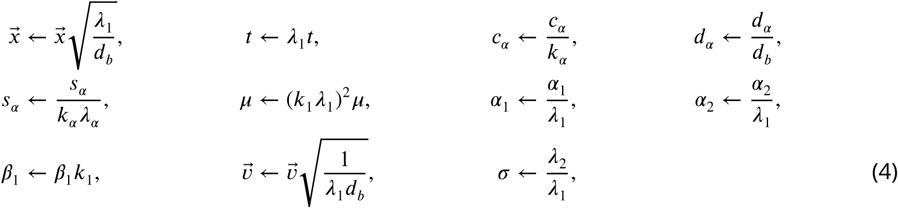

then our equations become,

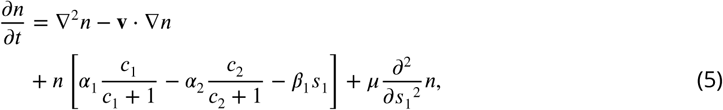

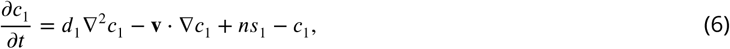

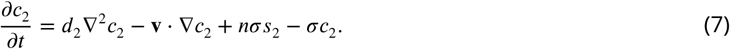

The analytical conclusions we derive from this system of equations has been guided and supplemented by an agent based stochastic simulation. Our simulation algorithm is as follows: microbes diffuse via a random walk, with step size derived from the diffusion constant and a bias dependent on the flow velocity. The microbes secrete chemicals locally that then diffuse and advect using a finite difference scheme. The microbes then reproduce or die with a probability dependent on their local fitness given by 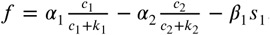. Upon reproduction, random mutations may alter the secretion rate of the public good –and thus the reproduction rate– of the microbes. The secretion rate is assumed to be heritable, and constant in time.

A summary of the system parameters is given in table 1, along with typical ranges for their values used in the simulations. The relevant ratios of parameters are consistent with those observed experimentally ***Kim*** (***1996***); ***Ma et al.*** (***2005***).

**Table 1.** Summary of system parameters.

| Parameter | Definition | Values |
| --- | --- | --- |
| $d_b$ | Microbial diffusion constant | $0.3906 \times 10^{-6} \text{ cm}^2 \text{ s}^{-1}$ |
| $d_1$ | Public good diffusion constant | $(1 \text{ to } 60) \times 10^{-6} \text{ cm}^2 \text{ s}^{-1}$ |
| $d_2$ | Waste diffusion constant | $(1 \text{ to } 50) \times 10^{-6} \text{ cm}^2 \text{ s}^{-1}$ |
| $v$ | Flow velocity | $(0 \text{ to } 100) \times 10^{-5} \text{ cm s}^{-1}$ |
| $\lambda_1$ | Public good decay constant | $5.0 \times 10^{-3} \text{ s}^{-1}$ |
| $\lambda_2$ | Waste decay constant | $1.5 \times 10^{-3} \text{ s}^{-1}$ |
| $k_1$ | Public good saturation | 0.01 |
| $k_2$ | Waste saturation | 0.10 |
| $s_1$ | Public good secretion rate | $(0 \text{ to } 1.0) \text{ s}^{-1}$ |
| $s_2$ | Waste secretion rate | $(0 \text{ to } 1.0) \text{ s}^{-1}$ |
| $\alpha_1$ | Benefit of public good | $7.5 \times 10^{-3} \text{ s}^{-1}$ |
| $\alpha_2$ | Harm of waste compound | $8.0 \times 10^{-3} \text{ s}^{-1}$ |
| $\beta_1$ | Cost of secretion | 0.2 |
| $\mu$ | Mutation rate | $(6.0 \text{ to } 20.0) \times 10^{-7} \text{ s}^{-1}$ |

### Social groups as Turing patterns

Diffusion can cause an instability that leads to the formation of intriguing patterns ***Turing*** (***1990***), which among other fields, have been investigated in ecological context ***Tian et al.*** (***2011***); ***Camara*** (***2011***); ***Baurmann et al.*** (***2007***). These so called Turing patterns typically form when an inhibiting agent has a diffusion length greater than that of an activating agent. For our model system, the waste compound and public good play the role of inhibiting and activating agents, and patterns manifest as cooperating microbial clusters (Fig 2).

**Figure 2.**
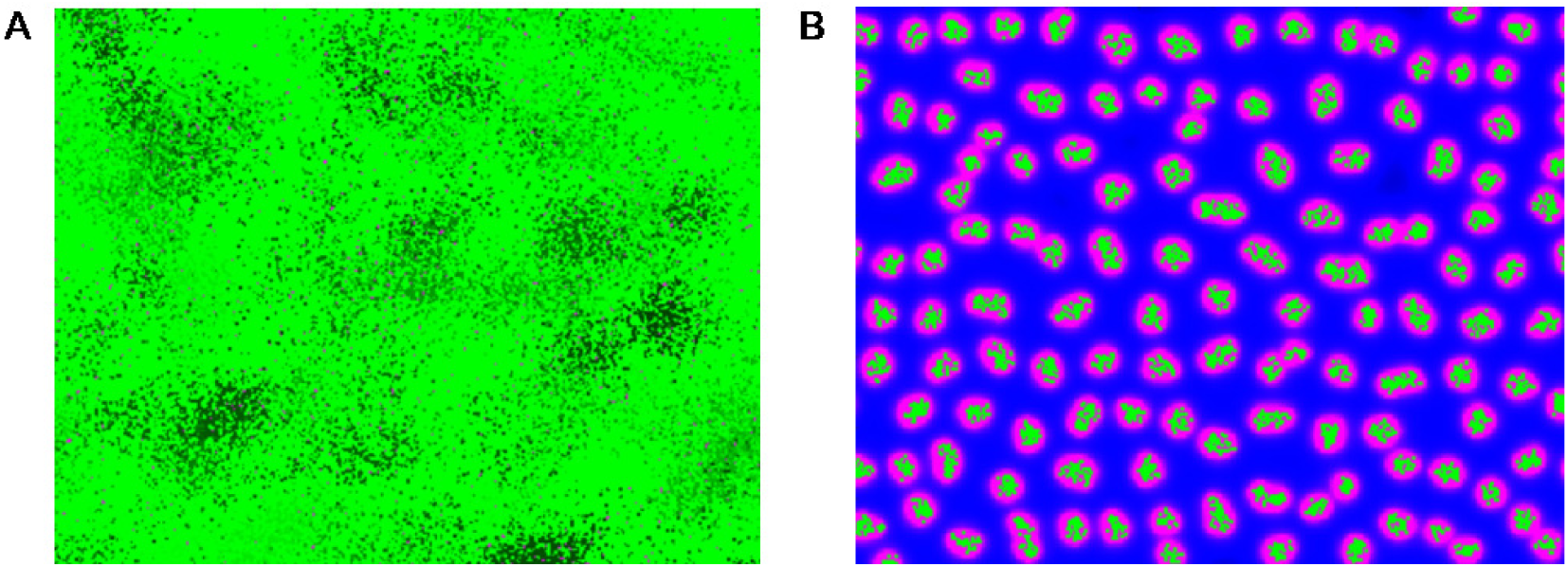
Homogeneous and group phases of microbes. Microbes (bright green dots) interact by secreting diffusive chemicals into their environment. The waste compound is shown as blue and the public good as red, the two combined is seen as magenta. **(A)** In the homogeneous phase the microbes spread to fill the domain.**(B)** In the group phase, when the diffusion length of the waste compound is larger an the diffusion length of public good, microbes form stable groups. As the microbes increase in number, the groups split apart and form new groups.

#### Steady states and linear stability

We first obtain the steady states in the absence of diffusion and investigate the stability of the system. The steady states are given by setting the reaction terms to zero,

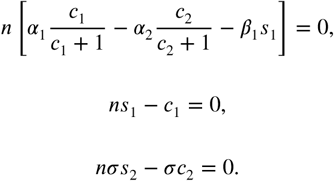

This gives, either the trivial solution 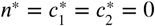, or the solutions:

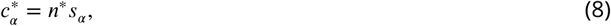

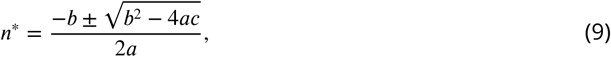

where

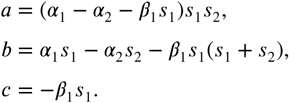

For this to be a sensible solution, we require *n*\* to be real and positive. This also imposes conditions on the system parameters.

Next, we establish the local stability of this solution by perturbing the system away from the steady state and expand up to first order. We take the perturbation 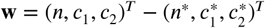, and substitute it into our system to get the linear system

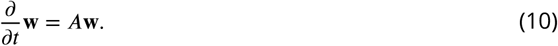

With the stability matrix, *A*, given as,

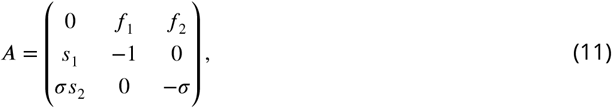

where

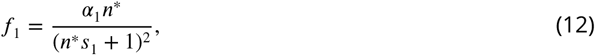

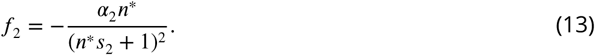

The system is stable if the eigenvalues, A of this matrix have a negative real part. The characteristic polynomial for the eigenvalues is given as Λ^3^ + *A*_0_ Λ^2^ + *B*_0_A + *C*_0_ = 0, where

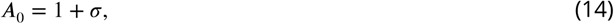

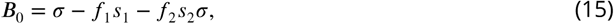

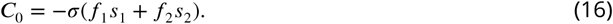

Since the first two coefficients of the characteristic equation are positive, by Descartes’ rule of signs, in order to get only negative real part eigenvalues, we need *B*_0_ and *C*_0_ to be positive as well. This is a requirement for linear stability. Thus, we have the conditions

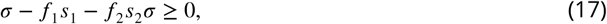

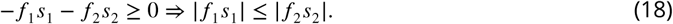

#### Diffusion driven instability

Next we include diffusion and analyze the instability caused by diffusion. Fourier expanding the solution

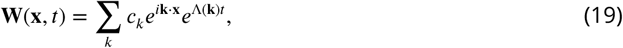

and plugging this into our equations, we get the eigenvalue equation, (−*k*^2^*D* + *A*)**W** = Λ**W**, where

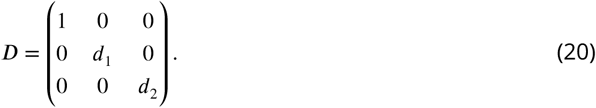

Solving for the eigenvalues gives a characteristic equation of the form Λ^3^ + *A* Λ^2^ + *B* Λ + *C* = 0, where

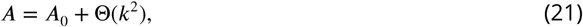

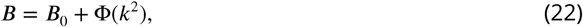

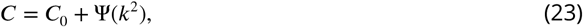

and the *k*^2^ dependent functions are,

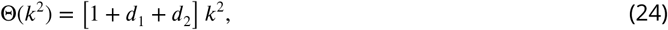

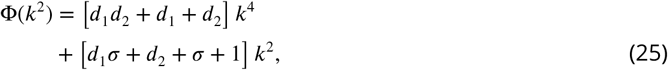

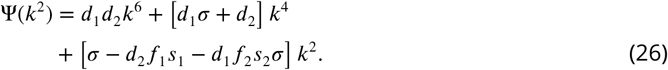

We can again use Descartes’ rule of signs, this time looking for an instability, which will happen when ψ is sufficiently negative. To be precise, the range of unstable wavenumbers satisfy,

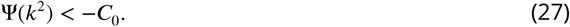

At the critical values for the diffusion parameters, the function ψ (*k*^2^) + *C*_0_ only vanishes at one point, the local minumum of ψ (*k*^2^). This occurs at the critical *k*^2^ value where d ψ/d*k*^2^ = 0, giving

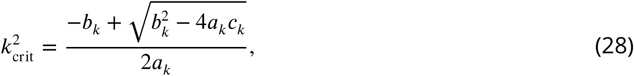

where

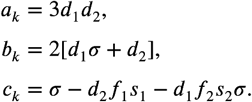

We take the positive root in (28) corresponding to the physical, positive *k*^2^.

For all combination of parameters giving rise to stable groups, we observe in our simulations that the group size approximated well by 2π/*k*_fast_, where, *k*_fast_ is the wave number corresponding to the fastest growing mode (i.e. the value that maximizes ψ (*k*^2^)). The size of the microbial clusters, as obtained by analytical theory 2π/*k*_fast_ and stochastic simulations are shown in the top row of Fig. 3. The color indicates the size of microbial groups.

**Figure 3.**
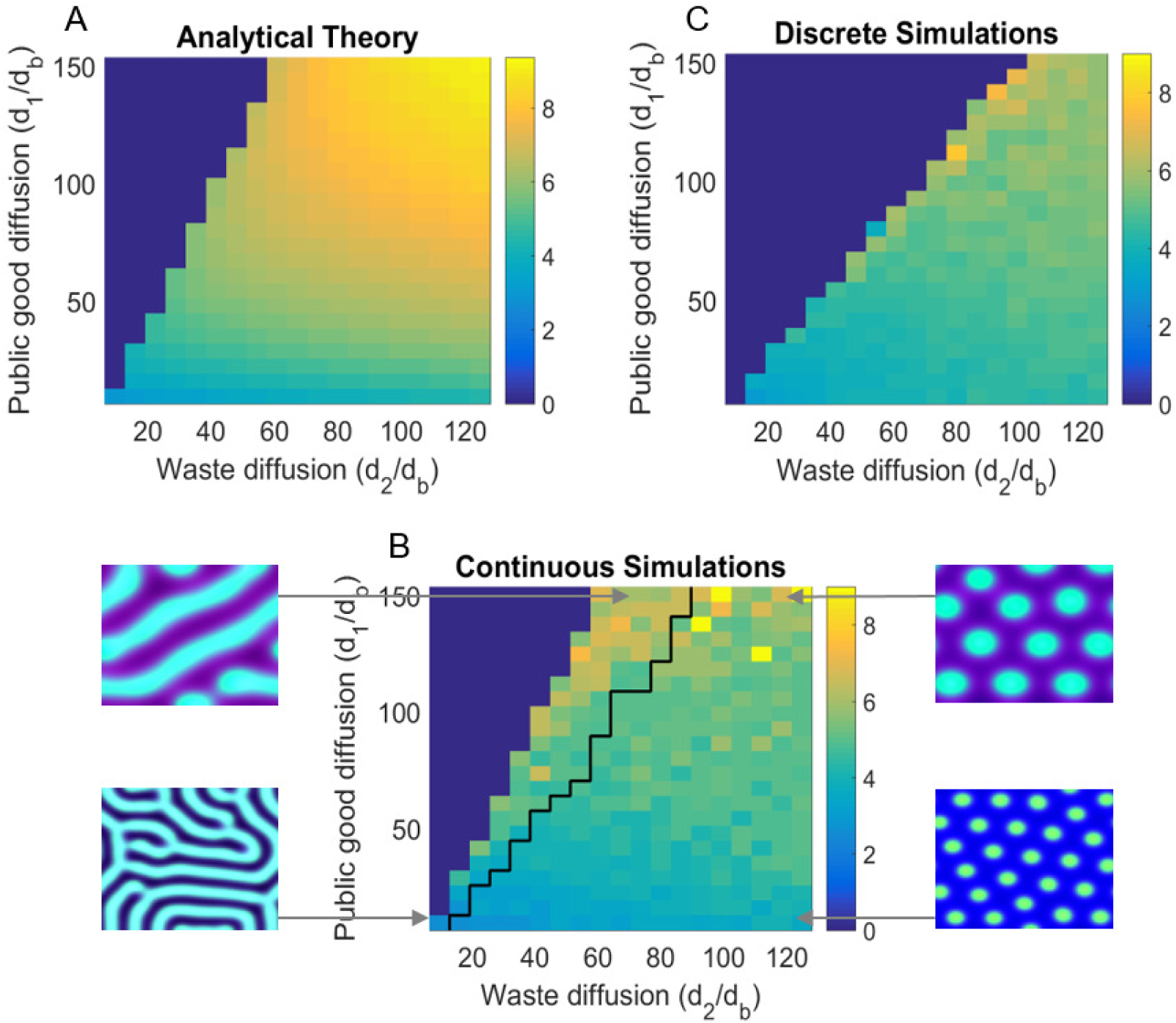
Turing analysis results. The top-left figure, **(A)** shows the group size 2π/*k*_fast_ as obtained by our theoretical analysis; whereas the bottom figure, **(B)** shows the same for continuous simulations, and the top-right figure, **(C)** is for agent based simulations. The black line in **(B)** divides parameters that give rise to striped patterns, and those that give rise to spots, corresponding images are shown to the left and right of **(B)**. Due to the discreteness of the agent based simulations, Turing patterns are not always stable where they might be in the continuous analogue. We see that the discrete simulations cut off around where we would see stripes in the continuous case, and do not see striped patterns in the discrete case. For different sets of parameters, we can also see Turing patterns in the discrete-stochastic case where they might not occur in the continuous case. For the region where Turing patterns are stable, the continuous theory gives a good prediction of group size and group reproduction rate.

Roughly speaking, if we view a microbial cluster as the cause of perturbation at a nearby location, the Turing instability will manifest as group reproduction. We should strongly caution that as the instability proceeds, the system moves away from the initial unstable fixed point around which it was linearized, and thus the exponential dependence in Eq. (19) should eventually break down. Nevertheless, the eigenvalues in the exponents still provide us with an approximate estimation of the group reproduction rate, within a factor of two near the phase boundary.

### Effect of shear on groups

We now move on to the key finding of the present theoretical study. Specifically, we investigate the effect of different flow velocity profiles on the social evolution of the system. A constant fluid flow merely amounts to a change in reference frame, which of course, does not change the evolutionary fate of the population. However, the derivative of the velocity causes significant changes to the social structure, both spatially and temporally. Specifically, we find that large shear rate causes microbial groups to distort, fragment, which in turn facilitates group reproduction. To investigate this effect in detail, we ran simulations for three fluid velocity distributions: Couette flow, Hagen-Poiseuille flow, and Rankine vortex.

#### Evolution of sociality in constant shear

To see the effect of shear on social evolution, we introduced Couette flow to the microbial habitat.

In this case, the flow velocity takes the form

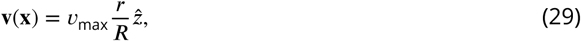

where *R* is the radius of the pipe, and 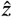 is the longitudinal direction.

The shear rate is the derivative of the flow velocity and is related to the maximum flow rate *v*_max_.

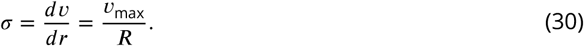

We ran simulations for various shear rates and diffusion constants and observed that shear does not significantly influence the *region* of parameter space that gives rise to cooperating groups. However *if* the system parameters are conducive to the formation of groups, shear tears groups apart and increases the rate at which spatially distinct cooperative clusters form.

We find that the reproduction rate *ω*(σ) of groups, depends linearly on the shear rate σ (Fig.4)

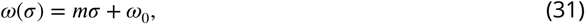

where ω_0_ is the reproduction rate solely due to microbial diffusion and can approximately be given by the Turing eigenvalue ω_0_ ≈ Λ_max_. The constant of proportionality *m* is given empirically from our simulations and depend on diffusion lengths and group density. This holds in the low density regime. Once the population density becomes large, group-group interactions slow the group reproduction rate and the population saturates.

We see that larger shear corresponds to a faster rate of group reproduction, thus enabling or enhancing social behavior in the microbial population.

We also find that the group population *N* increases linearly with shear,

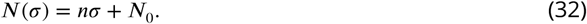

where *n* and *N*_0_ are the slope and intercept of the line in Fig.(4B).

Cooperation is stable if a group is able to fragment before a selfish strain emerges and proliferates in the group. Therefore, for stability, we need the take-over time to exceed the time it takes for a group to reproduce. A mutant emerges at a rate of *µN*(σ). Once a mutant emerges, it takes some time τ_*d*_ to spread to where the daughter group forms. The diffusion time *τ*_*d*_ will depend on where the mutant first emerges. Assuming a uniform distribution, and taking the diffusion time in two dimensions as a function of radius *r*, τ_*d*_ (*r*) = *r*^2^/4*d*_*b*_, we get an average diffusion time,

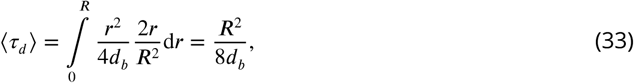

where *R* is the group radius. The take-over rate is then given by taking the inverse of the total take-over time, and the critical shear rate is given by equating the take-over rate with the reproduction rate,

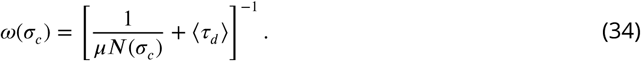

The critical shear rate σ_*c*_ above which the system can maintain stable cooperation is then given by the positive root,

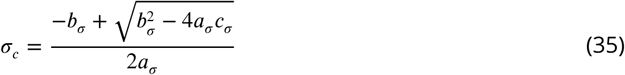

where

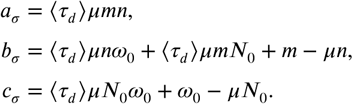

Eq. (35) is indicated by the vertical dashed lines in Fig 4 and agrees with the computationally observed critical shear reasonably well. It may be possible to improve this formula further by taking into account additional factors, such as the non-uniform spatial distribution of population within a group and the elongation of groups with shear. Furthermore, as the mutants increase in numbers it becomes more likely that one of them crosses over the the daughter, thereby reducing further the expected ⟨ τ_*d*_ ⟩. We see better agreement with analytical theory and simulations at lower mutation rates, (Fig 4), since these corrections are mainly to the diffusion time ⟨ τ_*d*_ ⟩., and become more significant at higher mutation rates, where ⟨ τ_*d*_ ⟩ ≫ 1/ μ*N*, (cf. Eq. (34)).

**Figure 4.**
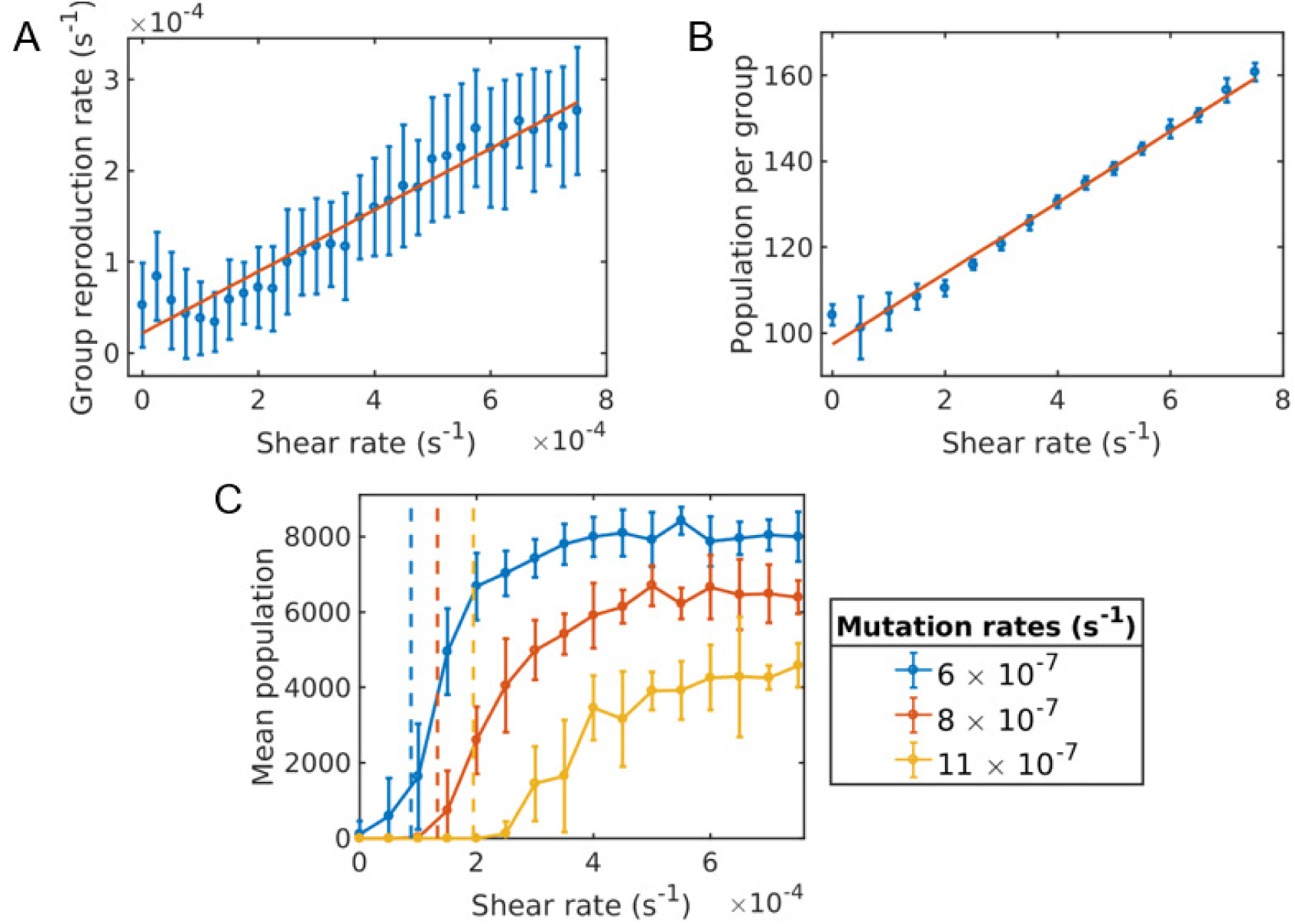
The effect of shear on microbial groups. **(A)** Group reproduction rate versus shear rate. We see that the group reproduction rate increases linearly with the shear rate. **(B)** Group population versus shear rate. As the shear distorts and elongates the group, the average group population also increases linearly with shear.**(c)** Average population versus shear rate for different mutation rates. Simulations were run for a time of 5.0 × 10^5^ s and averaged over 10 runs for each shear rate and mutation rate. The population goes extinct under larger mutation rates unless the shear rate is above the critical value. The critical shear values for different mutation rates are roughly obtained by Eq. (35) and are shown by the vertical dashed lines corresponding to curves of the same color.

If shear is below the critical value (35), the system will be in a non-social state. Ultimately, cheaters will take over, and wipe out all groups. When shear is increased above the critical value however, the system will transition to a stable social state, thereby maintaining its fitness and dense population indefinitely. Fig. 4 shows the long-time population of the system versus the shear rate. The population goes extinct under larger mutation rates unless the shear rate is above the critical value.

To summarize, the social state of the population can transition from a non-cooperative state to a cooperative one with increased flow shear rate.

#### Evolution of sociality in a flowing pipe

We now further generalize our results by looking at laminar flow with fixed boundaries. For Hagen-Poiseuille flow, the shear rate varies linearly with the radius, taking its maximum value adjacent to the boundaries, when *r* = *R*.

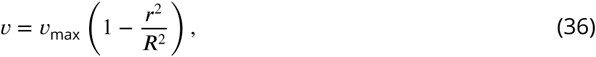

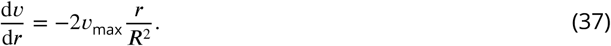

We therefore expect to see groups reproduce quicker at the boundary, leading to larger cooperation, higher average secretion rate, and larger population, which is indeed what we do see (Fig. 5).

**Figure 5.**
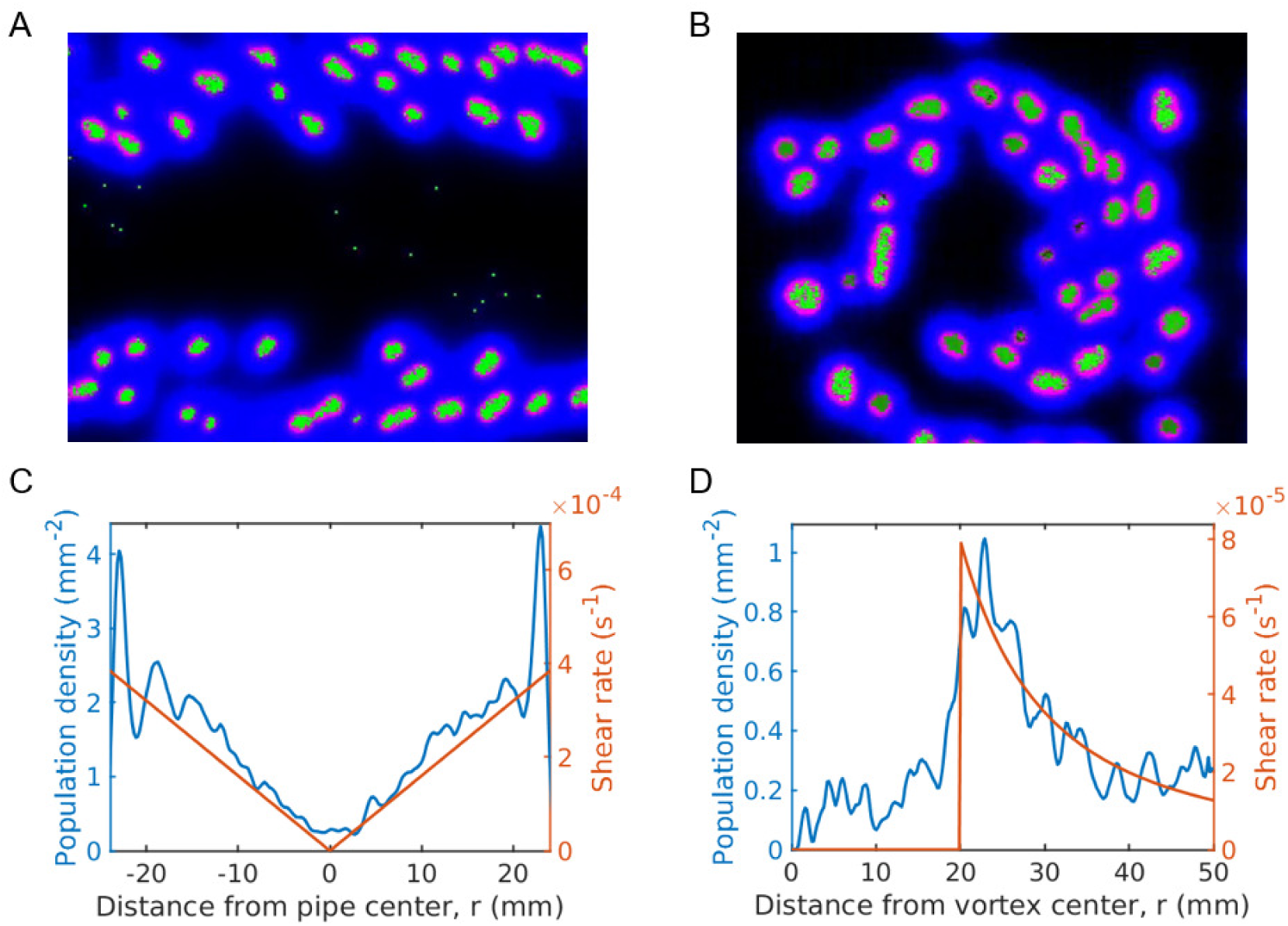
Average microbial population in pipe and vortex geometries. The top row gives simulation snapshots of the system in a Hagen-Poiseuille flow in a pipe **(A)** and of the system in a Rankine vortex **(B)**. The bottom row gives the average microbial population and the shear rate magnitude versus distance from the center of the pipe **(C)** and the center of the Rankine vortex **(D)**. Simulations were run for a duration of 2.0 × 10^5^ s under a mutation rate of μ = 2 × 10^-6^ s^-1^ and data was averaged over 100 runs. For Hagen-Poiseuille flow, we see that the population is larger at the boundaries, where the shear is also larger **(C)**. This is because groups reproduce quicker at the boundaries and are able to overcome take-over by mutation, whereas near the center they cannot. For the Rankine vortex we also see that the population follows very closely to the shear **(D)**, which suggests that the growth is proportional to shear. We caution that this holds in the low density limit. At higher densities the population saturates and is no longer proportional to shear. The undulations observed in the population plots are due to the finite size of the groups. Groups form layers of width equal to the group diameter. The population curve therefore shows undulations of width equal to the group width.

Earlier studies have proposed and shown that shear trapping due to the interaction between bacterial motility and fluid shear can result in preferential attachment to surfaces, ***Rusconi et al.*** (***2014***); ***Berke et al.*** (***2008***); ***Li et al.*** (***2011***). In a similar spirit, we suggest that inhabiting surfaces may have the additional advantage of enhanced sociality, due to shear driven group reproduction.

#### Evolution of sociality in vortices

In a vortex, the region above the critical shear value constitutes an annulus. Thus, we expect social behavior to be localized. Any point in the fluid outside this annulus will be taken over and destroyed by cheaters. In our simulations, at steady state we indeed see clusters whirling around exclusively within annulus, neither too near, nor too far from the vortex core. Life cannot exist outside this annulus, as cheaters kill these groups.

The Rankine vortex in two dimensions is characterized by a vortex radius *R* and a rotation rate Г. The velocity profile is given as,

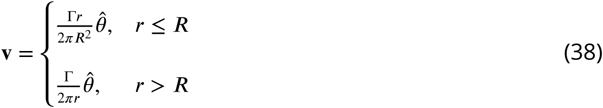

where *r*^2^ = *x*^2^ + *y*^2^.

The shear rate acting on a group acts tangential to the flow with a magnitude given by

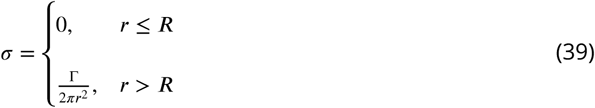

The shear rate is then a maximum at the minimum value of *r* which occurs at the vortex radius *R*. We therefore expect to see the largest concentration of groups at the vortex radius, which is what we observe in our simulations (Fig. 5).

## Limitations

While we paid close attention to physical realism, we also made important simplifying assumptions which under certain circumstances, may lead to incorrect conclusions. We caution the reader by enumerating the limitations of our model. First, since many microorganisms live in a low Reynolds number environment, we have chosen to neglect the inertia of microorganisms. However in reality, the microorganisms influence the flow around them. This effect will be particularly significant for a dense microbial population, especially when the microbes stick onto one other, or integrate via extracellular polymers. A more sophisticated model would include the coupling of the microbes to the flow. Secondly, the finite size and shapes of the microorganisms have been neglected. Instead, we have treated microbes as point particles, which will also invalidate our model in the dense population limit. Lastly, real microbes display a large number of complex behaviors such as biofilm formation and chemotactic migration. Here we have ignored the active response of microorganisms to the chemical gradients that surround them and to the surfaces they might attach and migrate. Instead, we took them as simple Brownian particles.

## Discussion

It is well known that spatial structure is crucial in the evolution of cooperation, ***Wilson et al.*** (***1992***); ***Taylor*** (***1992***); ***Lion and Baalen*** (***2008***). Many of these studies introduce these mechanisms “manually”, e.g. density regulation and migration are enforced by applying carrying capacities and migration rates to groups. In this study we have distanced ourselves from the typical game theoretic abstractions used to investigate evolution of cooperation. Instead we adopted a mechanical point of view. We investigated in detail, the fluid dynamical forces between microbes and their secretions, to understand how cooperation evolves among a population of planktonic microbes inhabiting in a flowing medium. In our first principles model, the spatial structuring and dispersion occur naturally from the physical dynamics.

We found that under certain conditions, microbes naturally form social communities, which then procreate new social communities of the same structure. More importantly, we discovered that regions of a fluid with large shear enable and promote the formation of such social groups. The mechanism behind this effect is that fluid shear distorts and tears apart microbial clusters, thereby limiting the spread of cheating mutants. Our proposed mechanism can be seen enhanced budding dispersal induced by shear flow.

Furthermore, while many theoretical ***Wilson et al.*** (***1992***); ***Taylor*** (***1992***) and experimental ***Küm-merli et al.*** (***2009b***) works have shown a viscous population is beneficial for cooperation, in the particular system we study, we find that the opposite is true. While Hamilton’s original argument is intuitive, we find that lower viscosity can introduce non-trivial fluid dynamical effects that alter the spatial structure of groups and counteract the spread of cheaters.

In our investigation, we found only certain regions of the fluid domain admits life, social or otherwise, as governed by the domain geometry and flow rate. From this perspective, it appears that evolution of sociality is a mechanical phenomenon.

In our physics-based model, groups with cheaters are negatively selected, and give way to those without cheaters. Furthermore, since the characteristics (e.g. shape, size, stability) of the group appears to be heritable, placing shear-induced cooperation reported here under the umbrella of “group selection” does not seem inappropriate ***Abbot et al.*** (***2011***). On the other hand the ensemble of groups do not exhibit any variation in their propensity to progenerate cheaters, neither is such propensity heritable. Rather, the progeneration of cheating is a (non-genetic) *symptom* that inevitably manifests in every group that has been around long enough. In this sense, it might be appropriate to view the emergence and spread of cheaters in a microbial population as a phenomenon of “aging”, in the non-evolutionary and mechanical sense, that any system consisting of a large number of interdependent components will inevitably and with increasing likelihood, fall apart ***Vural et al.*** (***2014***).

## Acknowledgments

This material is based upon work supported by the Defense Advanced Research Projects Agency under Contract No. HR0011-16-C-0062.

